# Potential Early Rabies Infection Detected in Two Raccoon Cases by LN34 pan-lyssavirus real-time RT-PCR Assay in Pennsylvania

**DOI:** 10.1101/2021.02.01.429287

**Authors:** Lisa Dettinger, Crystal M. Gigante, Maria Strohecker, Melanie Seiders, Puja Patel, Lillian A. Orciari, Pamela Yager, James Lute, Annette Regec, Yu Li, Dongxiang Xia

## Abstract

During 2017 – 2019, the Pennsylvania Department of Health Bureau of Laboratories (PABOL) tested 6,855 animal samples for rabies using both the gold standard direct fluorescent antibody (DFA) test and LN34 pan-lyssavirus reverse transcriptase quantitative PCR (RT-qPCR). Two samples (0.03 %) were identified as LN34 RT-qPCR positive after failure to detect rabies virus antigen during initial DFA testing: an adult raccoon collected in 2017 and a juvenile raccoon collected in 2019. After the positive PCR result, additional tissues were collected and re-tested by DFA, where very sparse, disperse antigen was observed. Tissues from both animals were submitted to the Centers for Disease Control and Prevention (CDC) for confirmatory testing, and were confirmed positive. At both PABOL and CDC, rabies virus antigen and RNA levels were much lower than for a typical rabies case. In addition, rabies virus antigen and RNA levels were higher in brain stem and rostral spinal cord than cerebellum, hippocampus and cortex. Cross-contamination was ruled out in the case of the 2019 juvenile raccoon by sequencing, as nucleoprotein and glycoprotein gene sequences displayed >1% nucleotide differences to sequences from all positive samples processed at PABOL within two weeks of the juvenile raccoon. Taken together, the low level of rabies virus in the central nervous system combined with presence in more caudal brain structures suggest the possibility of an early infection in both cases. These two cases highlight the increased sensitivity and ease of interpretation of LN34 RT-qPCR in rabies diagnostics for the identification of low positive cases.

## Introduction

Rabies is a fatal but preventable infectious disease that causes approximately 60,000 human deaths worldwide each year (1). In the United States rabies causes few human deaths thanks to the elimination of rabies variants maintained in domestic dogs and large-scale, sustained rabies control efforts (2–4). Still, rabies endemic in wildlife presents a threat to humans and domestic animals. Rabies surveillance in the United States involves over 125 rabies testing laboratories (5). Each year, more than 100,000 animal samples are tested, and approximately 5,000 rabid animals are identified (5, 6). The major reservoirs are bats, raccoons, skunks, and foxes. Several distinct rabies virus variants are endemic in the United States; these variants are named based on the known or presumptive reservoir species associated with enzootic transmission. Rabies is maintained in many species of bats across the continent, and several bat rabies virus variants have been identified (7–10). Rabies variants endemic in raccoons, skunks, and foxes have distinct geographic distributions with few areas of overlap (5, 6). A single variant known as “Eastern Raccoon” rabies virus variant is endemic in raccoons along the East Coast.

In the United States, rabies diagnostic testing is predominantly performed using the international gold standard Direct Fluorescent Antibody test (DFA). DFA has been a reliable and sensitive rabies diagnostic test for over 60 years; however, there is a need to assess newer methods. The World Health Organization (WHO) and World Organization for Animal Health (OIE) recognize reverse transcriptase polymerase chain reaction (RT-PCR) as a diagnostic test for the detection of rabies virus (11, 12). Molecular methods such as reverse transcriptase quantitative PCR (RT-qPCR) provide several advantages over DFA testing. Many public health laboratories routinely perform RT-qPCR for detection of other pathogens and already have the equipment and expertise to implement a rabies RT-qPCR test. DFA, however, requires fluorescence microscopy expertise, which is less and less frequently used in diagnosis of other pathogens.

RT-PCR is not currently recommended for primary diagnostic testing of rabies samples in the United States, though it can be used as a confirmatory test (13). Laboratories across the country are currently implementing the LN34 RT-qPCR assay (14, 15) for confirmatory rabies testing. The Pennsylvania Department of Health Bureau of Laboratories (PABOL) routinely tests every suspect rabies sample by DFA and LN34 RT-qPCR. PABOL tests approximately 2,800 animals associated with human exposures for rabies annually. On average, 114 positive rabies cases are identified (4%). Raccoons and bats are the major reservoirs in Pennsylvania, and Eastern Raccoon rabies virus variant and several bat variants are endemic.

During 2017 – 2019, 6,855 animals were tested by both DFA and PCR at PABOL. Of those tested, only two (0.03%) were initially DFA negative but positive by LN34 RT-qPCR. In June 2017, an adult raccoon in Carbon County, PA, attacked a chicken and charged an individual. On June 1^st^ the animal was euthanized and submitted for rabies testing. Initial DFA test was determined to be negative, but rabies virus RNA was detected by LN34 RT-qPCR. In June 2019, a mother raccoon was hit and killed on a road in Venango County, PA, leaving behind two young offspring. The two juvenile raccoons were taken into a home and kept from June 11^th^ to 12^th^, during which time they were handled by four persons. On June 13^th^, both juveniles were euthanized and submitted for rabies testing. One juvenile was negative by both DFA and LN34 RT-qPCR. The other juvenile tested positive by LN34 RT-qPCR after initial negative DFA result. The following report describes the subsequent investigation into these two cases.

## Materials and Methods

### Samples

Samples were submitted to PABOL as part of routine rabies surveillance and diagnostic testing. Animal collection was not performed as part of this study; therefore, institutional animal care and use committee approval was not necessary.

### Direct Fluorescent Antibody (DFA) Test

#### PABOL

Brain tissue representing a full transverse cross section of brain stem and three lobes of cerebellum and/or hippocampi were minced together. These brain tissue preparations were tested used a modification of the minimum United States national standard protocol (national standard protocol) (13). Additional details can be found in the Supplemental Text.

#### CDC

Samples were tested according to the Protocol for Postmortem Diagnosis of Rabies in Animals by Direct Fluorescent Antibody Testing, A Minimum Standard for Rabies Diagnosis in the United States and Direct Fluorescent Antibody Test, WHO, Laboratory Techniques in Rabies [13, 16]. Additional details can be found in the Supplemental Text.

### Real-time RT-PCR (RT-qPCR)

Tissue representing a full cross section of brain stem and all three lobes of cerebellum was transferred to TRIzol Reagent (Life Technologies 15596018) and then extracted using Direct-zol RNA MiniPrep kit (R2052 Zymo, Irvine, CA, USA) following the published protocol for LN34 RT-qPCR (14). Additional RT-qPCR testing of separate tissues was performed for brain stem, rostral spinal cord, cerebellum, hippocampi and cortex. Samples were tested in duplicate on the Applied Biosystems 7500 Fast Dx platform at PABOL. Samples were tested in triplicate on Applied Biosystems ViiA7 platform at CDC. LN34 Cq values were used to compare relative levels of viral RNA in different brain regions. CDC operators were not blinded to the samples.

Quantification of RT-qPCR results was performed using the delta delta Cq method (ΔΔCq or ddCq) (17). Average LN34 and beta actin Cq values were calculated for each brain region examined. Average actin Cq value was subtracted from the average LN34 Cq value for each brain region to calculate the ΔCq. Brain stem was chosen as the reference tissue, so ΔCq for the brain stem was subtracted from ΔCq for other brain regions to calculate ΔΔCq for each brain region. Amount of target was estimated as 1.93^−ΔΔCq^, based on the efficiency of the LN34 assay as 93% for rabies virus based on previous estimation (14). Plot was generated in RStudio (18) using ggplot2 (19) and finished in Inkscape 0.91 (inkscape.org).

### Sequencing

Rabies virus sequencing was performed at CDC for the 2019 juvenile raccoon case and four additional positive samples that were manipulated at PABOL within two weeks of the 2019 case to rule out potential contamination. Complete rabies virus nucleoprotein and glycoprotein gene sequences were generated from rabies virus RNA extracted using Direct-zol RNA MiniPrep kit (R2052 Zymo, Irvine, CA, USA). Complete nucleoprotein and glycoprotein genes were amplified using Takara long amplicon Taq polymerase with GC buffers (RR02AG Takara Bio USA, Mountain View, CA, USA) using the primers indicated in Table 1 after cDNA synthesis using random hexamer primers and Roche AMV reverse transcriptase (10109118001 Roche, Sigma-Aldrich, St. Louis, MO, USA). Samples were multiplexed using Takara long amplicon Taq polymerase with GC buffers following the manufacturer’s instructions for PCR barcoding for nanopore sequencing (EXP-PBC096 Oxford Nanopore Technologies, Oxford, UK). Samples were pooled and sequenced using the Oxford Nanopore MinION, following the manufacturer’s instructions for the ligation sequencing kit (SQK-LSK108 Oxford Nanopore Technologies, Oxford, UK). Consensus sequences were generated in CLC Genomics Workbench 12 (Qiagen, Venlo, Netherlands) after read mapping to rabies virus reference genomes using bwa mem -x ont2d (Li arXiv:1303.3997v1 2013) and were polished using nanopolish version 0.6.0 (https://github.com/jts/nanopolish/). Manual indel correction was then performed as described previously for the coding regions of the nucleoprotein and glycoprotein genes (20)[Gigante in preparation]. Sequence differences were determined based on coding region alignments generated using mafft v7.308 (21, 22) in geneious 9.1.4 (Biomatters, Inc., Newark, NJ, USA). Phylogenetic analysis was performed by Maximum Likelihood in Mega 7.0.26 (23) using GTR+G+I model of evolution, which was determined using model test in Mega7.

**Table 1.**
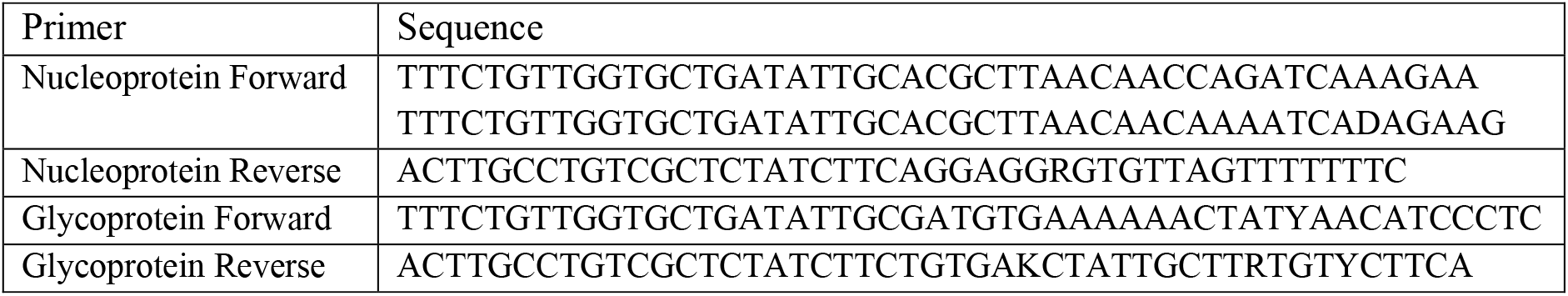
Rabies primers used for sequencing in this study. Note: primers include 5′ sequence for adding Oxford Nanopore barcode sequences by PCR.

### Data Availability

Sequences were deposited under GenBank accession numbers (awaiting accession numbers).

## Results

### PABOL DFA and PCR Testing

During 2017 – 2019, PABOL tested 6,855 animals submitted for rabies testing. A total of 342 were positive (4.06%). Raccoon was identified as the leading host species, with 123 rabid raccoons identified, followed by cats (91), foxes (45), bats (43) and skunks (17) (Figure 1).

**Figure 1.**
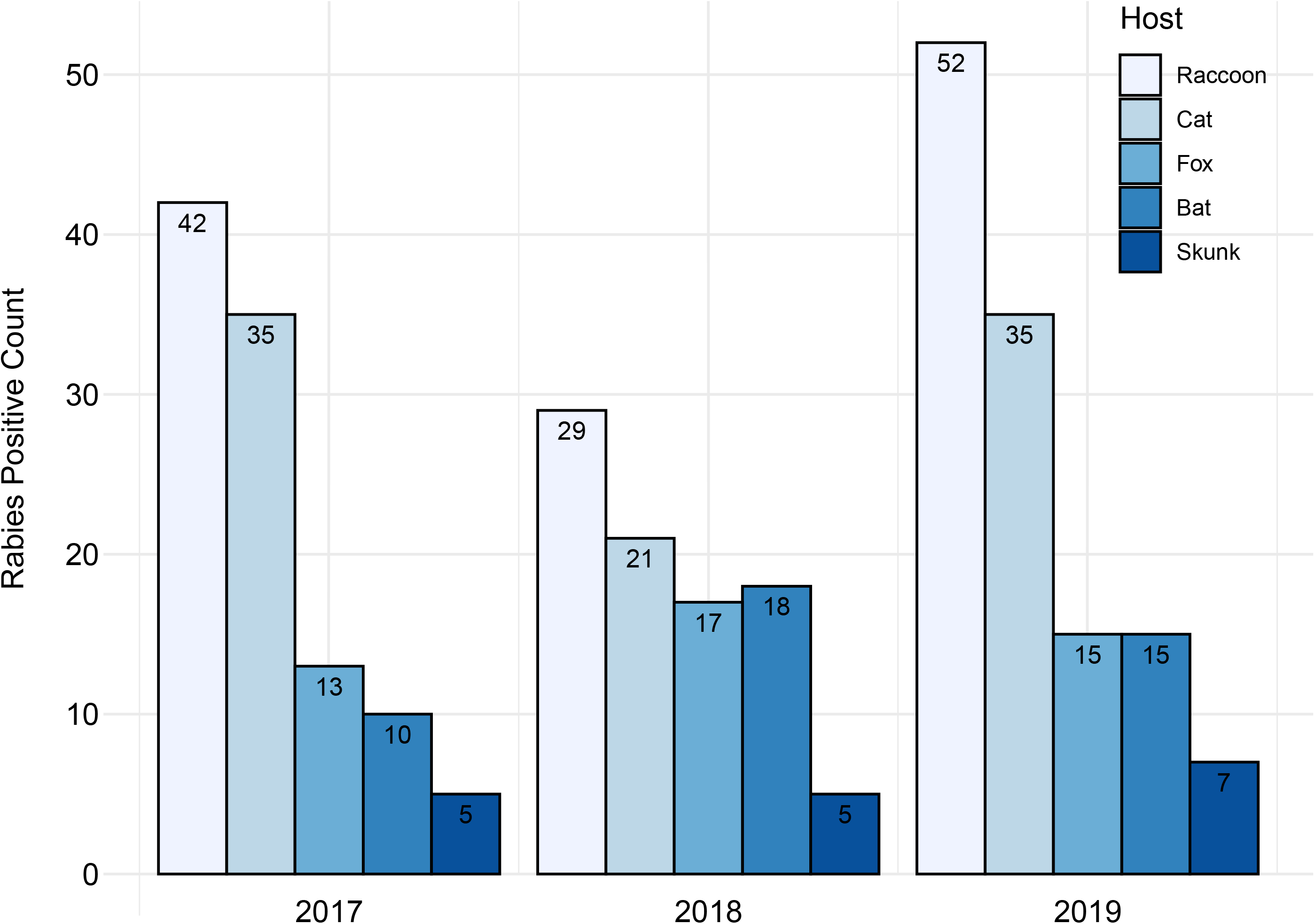
Distributions of positive rabies samples tested at PABOL during 2017 – 2019 by host animal. Raccoons accounted for 37% (42/113), 31% (29/95) and 39% (52/134) of positive cases each year, respectively.

Since 2018, PABOL routinely tests all rabies samples by DFA in parallel with LN34 RT-qPCR. PABOL participated in a LN34 RT-qPCR pilot study with the Centers for Disease Control and Prevention (CDC) (14) in 2016 and fully implemented PCR along with DFA testing for all samples in 2018. Among 6,855 samples tested in 2017 – 2019, discordant results were identified for only two cases (0.03%): an adult raccoon tested in 2017 (sample 1130) and a juvenile raccoon tested in 2019 (sample 1059). In these two cases, the initial DFA tests were negative for rabies antigen; however, rabies virus RNA was detected by LN34 RT-qPCR (Table 2).

**Table 2.**
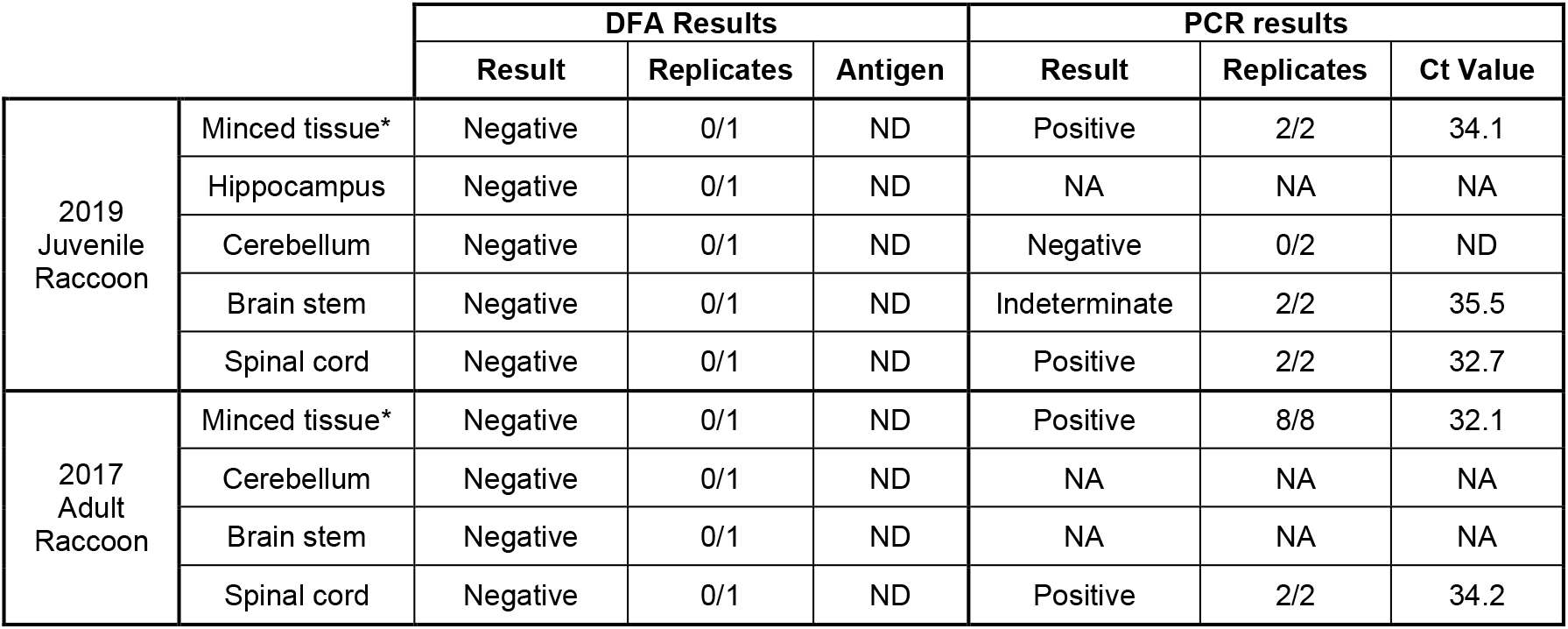
DFA and RT-qPCR results from PABOL. Average rabies virus (LN34) Cq value is given for each sample, where lower Cq value indicates higher rabies virus RNA level. Cq value >35 was used to define indeterminate result for LN34 RT-qPCR, based on previous publication [15]. NA: sample not available. ND: not detected. *Initial minced tissue from hippocampus, cerebellum, brain stem and spinal cord.

In both cases, the original tissues were reprocessed, taking separate samples from different regions of the brain, including rostral spinal cord, brain stem, cerebellum, and hippocampus. These separate brain tissues were tested by both DFA and LN34 RT-qPCR. Upon re-testing, some atypical, sparse staining was observed by DFA in brain stem and spinal cord impressions but was notably absent from cerebellum and hippocampus. Rabies virus RNA levels were low in all tissues tested for RT-qPCR, with the highest levels (lowest quantification cycle (Cq) values) in spinal cord and brain stem and the lowest levels in cerebellum and hippocampus (Table 2).

### CDC DFA and PCR Testing

Brain samples were sent to the Poxvirus and Rabies Branch at CDC for confirmatory testing by DFA and LN34 RT-qPCR. Aliquots of both samples were confirmed positive with low antigen distribution; however, antigen distribution varied in different regions of the brain (Table 3). For both samples, all impressions prepared from brain stem or rostral spinal cord tissue were positive, with typical antigen in <10% of fields examined. Cerebellum tissue also produced positive DFA results; however, typical rabies antigen was observed in only 2/6 slides for the 2017 adult raccoon and 3/5 slides for the 2019 juvenile raccoon. Rabies antigen distribution in the positive cerebellum slides was also in <10% of fields. Impressions from cortex and hippocampus were tested from the 2019 juvenile raccoon. One slide out of 6 showed atypical staining; the remaining 5 cortex/hippocampus slides did not contain typical rabies antigen, and the result was indeterminate.

**Table 3.** DFA and RT-qPCR results from CDC. Antigen distribution refers to percent of fields showing positive rabies antigen. Average rabies virus (LN34) Cq value is given for each sample, where lower Cq value indicates higher rabies virus RNA level. Cq value >35 was used to define indeterminate result for LN34 RT-qPCR, based on previous publication (14). NA sample not tested. *Average Cq values do not include replicates that did not produce Cq values (1/3 for cortex/hippocampus and 2/6 for cerebellum). The 2019 cortex/hippocampus tissue also contained remaining brain tissue from the head.

|  |  | DFA Results |  |  | PCR results |  |  |
| --- | --- | --- | --- | --- | --- | --- | --- |
|  |  | Result | Replicates | Antigen | Result | Replicates | Ct Value |
| 2019 Juvenile Raccoon | Cortex/Hippocampus | Indeterminate | 1/6 | Atypical | Indeterminate | 2/3 | 42.6* |
|  | Cerebellum | Positive | 3/5 | <10% | Positive | 1/6 | 39.0* |
|  | Brain stem | Positive | 3/3 | <10% | Positive | 3/3 | 35.0 |
|  | Spinal cord | NA | NA | NA | Positive | 3/3 | 32.4 |
| 2017 Adult Raccoon | Cerebellum | Positive | 2/6 | <10% | Positive | 3/3 | 31.8 |
|  | Brain stem | Positive | 1/1 | <10% | Positive | 3/3 | 31.4 |
|  | Spinal cord | Positive | 1/1 | <10% | NA | NA | NA |

In addition to brain tissues, tissue homogenates in TRIzol and extracted RNA were submitted to CDC for RT-qPCR testing for the 2019 juvenile raccoon case. All aliquots used for DFA brain impressions were tested by RT-qPCR, but TRIzol or RNA samples could not be tested by DFA. All samples exhibited amplification, indicating the presence of rabies virus RNA; although, in some cases, amplification did not reach the threshold or the Cq value was later than the cut-off for a positive sample, indicating an indeterminate result. Samples taken from brain stem, cerebellum or rostral spinal cord were all positive by RT-qPCR. All replicates produced positive results for brain stem and spinal cord samples and the cerebellum tissue from the 2017 case (Table 3). For the 2019 juvenile raccoon, only 1 out of 6 replicates from two cerebellum samples produced Cq value <35 required for a positive result. Cortex and hippocampus tissue from the 2019 juvenile raccoon produced an indeterminate result because Cq values were ≥35 (14). Rabies virus RNA levels were highest in the spinal cord and brain stem and lowest in the cortex/hippocampus (Figures 2 and 3).

**Figure 2.**
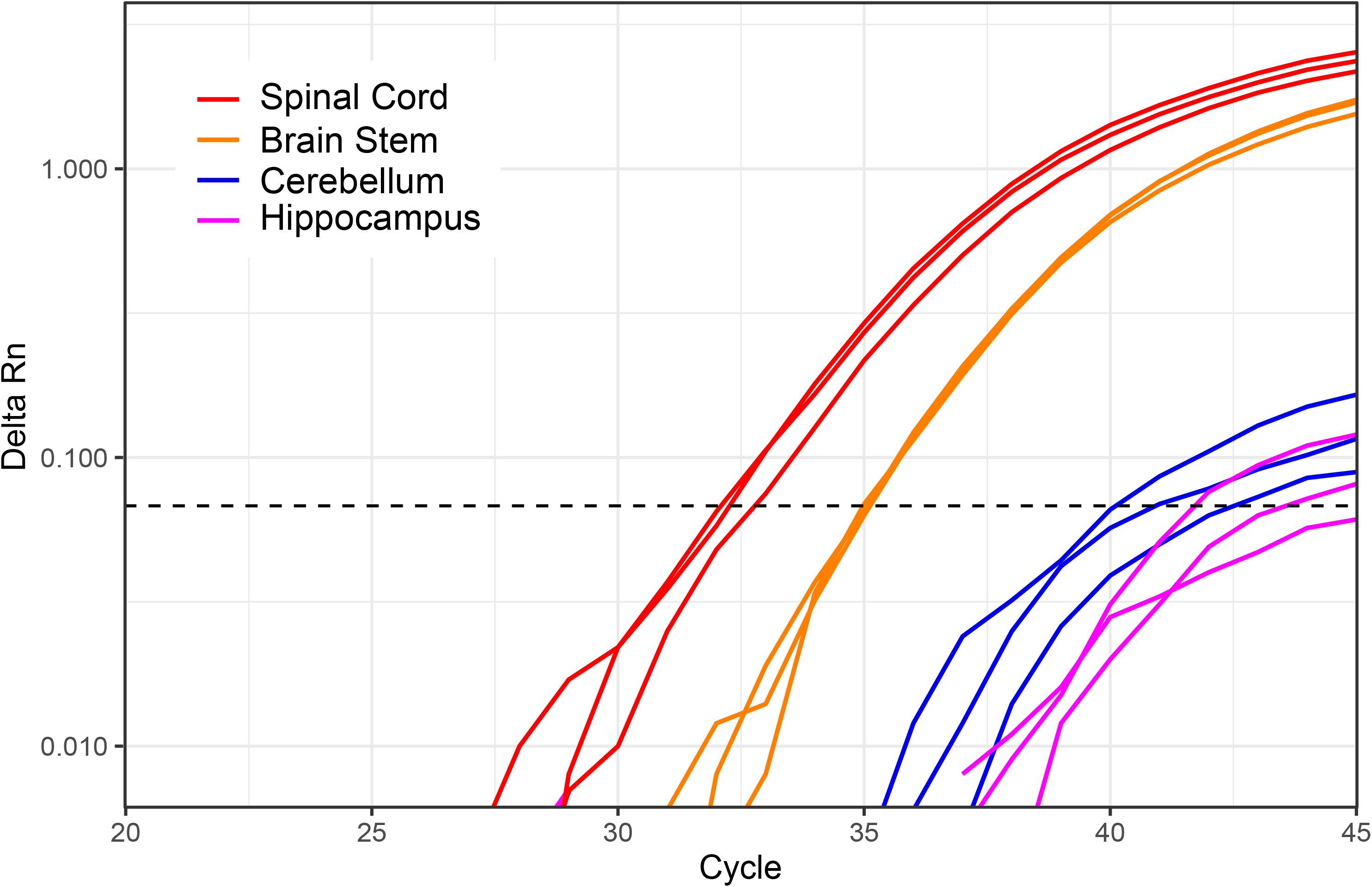
PCR amplification curves produced by LN34 RT-qPCR of different brain tissues from the 2019 PA juvenile raccoon. Increasing rabies virus RNA level, as indicated by earlier amplification, can be observed from hippocampus to cerebellum to brain stem to rostral spinal cord. Threshold used for Cq value calculation is shown by dotted line. Triplicate results are shown.

**Figure 3.**
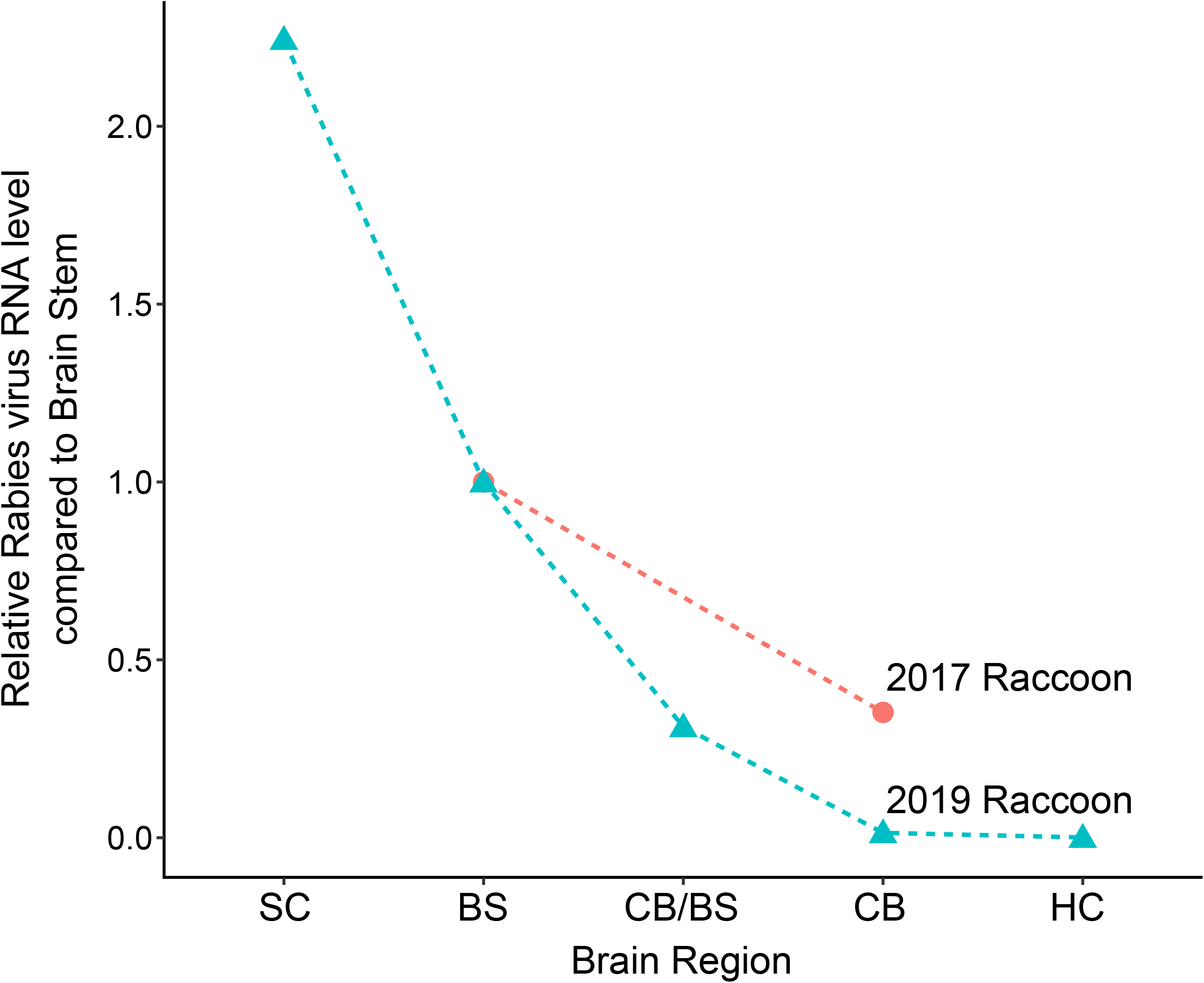
Relative rabies virus RNA level in different brain regions of 2017 and 2019 PA raccoon samples. Rabies virus RNA (LN34 Cq value) was normalized to beta actin level and compared to brain stem using the ΔΔCq method (17). SC spinal cord, BS brain stem, CB/BS mix of brain stem and cerebellum, CB cerebellum, HC hippocampus/cortex.

### Investigation into potential cross-contamination

Taken together, the low rabies virus RNA level and distribution pattern of antigen and RNA (highest in caudal brain regions and lowest in rostral regions) in these two cases could indicate early infection or cross-contamination. To rule-out the possibility of cross-contamination, rabies virus sequencing was performed on the 2019 juvenile case and all positive samples processed at PABOL within two weeks. These included grey fox sample 997 (processed 6/11), bat sample 1018 (processed 6/13), grey fox sample 1090 (processed 6/19), and cat sample 846 (used as a positive control the week juvenile raccoon 1059 was tested). Sequencing was not performed for the 2017 case because samples were no longer available.

Complete nucleoprotein and glycoprotein gene sequences were generated and compared to publicly available reference sequences from representative rabies virus variants. BLAST search of rabies virus sequences from the 2019 juvenile raccoon revealed > 99% nucleotide identity with Eastern Raccoon rabies virus variant isolates from the eastern US. Phylogenetic analysis revealed the 2019 juvenile raccoon sequence clustered with other Eastern Raccoon variant sequences from PA and reference sequence MK540681 (raccoon from NY 1991) (Figure 4). PA cat 846, PA fox 997 and PA fox 1090 clustered with the 2019 juvenile raccoon sequence within with the Eastern Raccoon variant clade. PA bat 1018 clustered with reference JQ685920, collected from a big brown bat in PA in 1984 and rabies virus variant EF-E1 (24) that is maintained in the big brown bat, *Eptesicus fuscus,* in the eastern US.

**Figure 4.**
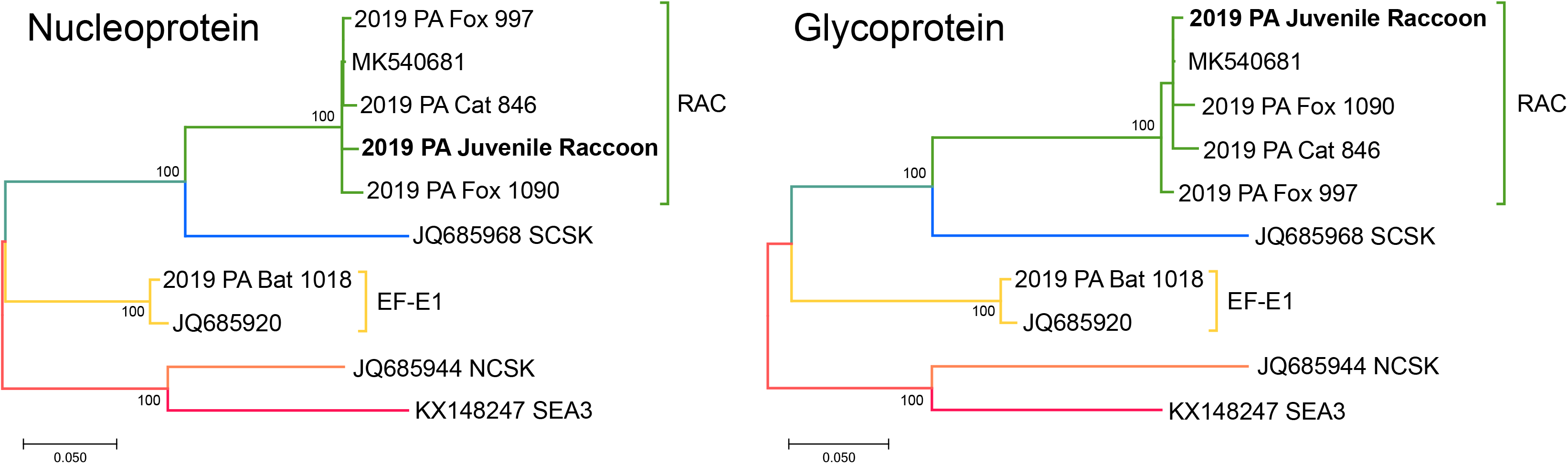
Phylogenetic trees showing clustering of PA 2019 juvenile raccoon rabies virus nucleoprotein (left) and glycoprotein (right) sequences with other rabies positive PA and reference sequences. Reference sequences from Eastern Raccoon (RAC), South Central Skunk (SCSK), *Eptesicus fuscus* Eastern 1 (EF-E1), North Central Skunk (NCSK) and South East Asia 3 (SEA3) rabies virus variants are shown with accession numbers. Branch color indicates variant: green is RAC, blue is SCSK, yellow is EF-E1, orange is NCSK and red is SEA3. The percentage of trees in which the associated taxa clustered together is shown next to the branches (based on 1,000 bootstraps). Scale bar indicates number of substitutions per site.

The 2019 juvenile raccoon sequences exhibited many differences from all other PABOL samples processed within two weeks (Table 4). The nucleoprotein gene had 17 – 23 nucleotide differences compared to the Eastern Raccoon variant samples and 195 differences compared to bat sample 1018. The glycoprotein gene had 23 – 26 changes relative to the Eastern Raccoon variant samples and 270 changes relative to the bat sample. The closest PABOL sequence was fox sample 997, which exhibited 98.7% and 98.5% identity to the nucleoprotein and glycoprotein genes, respectively. The 2019 juvenile raccoon sequences were more similar to an Eastern Raccoon variant isolate from NY in 1991 (MK540681, with 99.04% and 99.49% identity to nucleoprotein and glycoprotein genes, respectively). Taken together, these data suggest contamination was unlikely the cause of the positive PCR result.

**Table 4.** Distance matrix showing number of nucleotide differences between the rabies virus isolated from the 2019 juvenile raccoon (2019 PA Juv Rac), samples processed around the same time as the juvenile raccoon and reference sequences. Numbers in the top right are differences in the nucleoprotein coding region; numbers in the bottom left are differences in the glycoprotein coding region. Differences relative to the 2019 juvenile raccoon are shown in bold. Reference sequences from Eastern Raccoon (RAC), South Central Skunk (SCSK), *Eptesicus fuscus* Eastern 1 (EF-E1), and North Central Skunk (NCSK) rabies virus variants are shown with accession numbers. Color indicates rabies virus variant: green is RAC, blue is SCSK, yellow is EF-E1 and orange is NCSK.

|  | JQ685944<br>NCSK | 2019 PA<br>Cat 846 | 2019 PA<br>Fox 997 | MK540681<br>RAC | **2019 PA**<br>**Juv Rac** | 2019 PA<br>Fox 090 | JQ685968<br>SCSK | 2019 PA<br>Bat 018 | JQ685920<br>EF-E1 |
| --- | --- | --- | --- | --- | --- | --- | --- | --- | --- |
| JQ685944 NCSK |  | 230 | 228 | 231 | 234 | 235 | 239 | 190 | 191 |
| 2019 PA Cat 846 | 329 |  | 14 | 10 | **21** | 20 | 168 | 197 | 198 |
| 2019 PA Fox 997 | 335 | 37 |  | 6 | **17** | 18 | 165 | 194 | 195 |
| MK540681 RAC | 332 | 20 | 19 |  | 13 | 16 | 164 | 196 | 195 |
| **2019 PA Juv Rac** | 335 | **26** | **23** | 8 |  | **25** | 168 | **195** | 194 |
| 2019 PA Fox 090 | 331 | 31 | 32 | 17 | **23** |  | 164 | 200 | 198 |
| JQ685968 SCSK | 339 | 251 | 244 | 248 | 252 | 255 |  | 211 | 209 |
| 2019 PA Bat 018 | 291 | 268 | 273 | 268 | **270** | 264 | 285 |  | 18 |
| JQ685920 EF-E1 | 290 | 270 | 275 | 270 | 272 | 266 | 296 | 19 |  |

## Discussion

We describe two cases where LN34 RT-qPCR identified rabies cases with very low viral RNA after initial DFA testing failed to detect the presence of rabies virus antigen. Repeat testing at PABOL and confirmatory testing at CDC confirmed both as positive rabies cases, and appropriate public health response was initiated. These cases highlight the sensitivity and objectivity of PCR in cases with low rabies virus antigen and RNA and support the addition of PCR for routine rabies diagnostic testing.

### Real-time RT-PCR in rabies diagnosis of low positive samples

A false negative result for a rabies diagnostic test is extremely serious because rabies is nearly always fatal if post exposure treatment is not administered promptly. The DFA has been used for over 60 years in the United States with no known deaths caused by failures to detect rabies cases. With these two cases, RT-qPCR demonstrated higher sensitivity than DFA at PABOL, and the reasons behind this are worth considering.

PABOL tests thousands of samples each year, and the concordance rate for DFA with PCR was 99.97% for 6,855 samples. If there was a systemic issue with DFA testing at PABOL, a lower concordance rate with LN34 would be expected, similar to what has been reported previously for laboratories with systemic DFA issues (false positives) (14). One observation worth noting is the practice of making impressions from minced brain tissues at PABOL. The United States national standard protocol (13) and WHO (16) recommend that impressions are taken directly from tissue for DFA testing. However, repeat testing of tissue impressions from these cases also produced negative DFA results at PABOL. The national standard protocol (13) was developed to avoid differences between laboratories.

In general, any differences in DFA test procedures between laboratories can affect test results (25) and should be avoided. The DFA procedure could vary between laboratories due to differences in commercial monoclonal antibody reagents or if optimal working dilutions of conjugate were not prepared properly (13, 26). Differences in fluorescence microscopes and objective lens quality could, in theory, produce different results for a sample with extremely low antigen level. The DFA relies heavily on the expertise of the person interpreting results, who must be able to distinguish typical fluorescent rabies virus antigen from non-specific fluorescent objects such as bacteria or artifacts in the tissue. All atypical, weak or unusual tests are repeated using a specificity control or sent to CDC for confirmation.

During initial testing at PABOL by DFA, one slide containing brain stem and cerebellum was tested for each sample and results were negative. Prompted by the positive PCR result, DFA re-testing was initiated. Many impressions were made from different brain regions, increasing opportunity to detect sparse antigen at both labs. However, antigen was detected in every brain stem impression tested at CDC for both samples.

In contrast to DFA, PCR methods are easier to standardize, and result interpretation is inherently more objective. Primer and probe sequences and concentration can be defined in protocols for high reproducibility between laboratories and uniformity between manufacturers and lots. Currently, CDC provides a standardized positive control to ensure proper performance of the LN34 across laboratories. Test output is a quantitative Cq value, which determines positive, negative or indeterminate result based on its numeric value. However, the high sensitivity of PCR can lead to false positive results caused by cross-contamination, especially in laboratories inexperienced with PCR. In most cases, cross-contamination can be avoided through good laboratory practice.

Many laboratories in the United States already employ RT-PCR as a confirmatory test for rabies when DFA exhibits non-specific staining. In these cases, RT-PCR can confirm a negative result and avoid unnecessary post-exposure prophylaxis for exposed humans or reduce quarantine for exposed animals. The findings from the two PA raccoon cases support expanding the role of PCR in rabies diagnosis in the United States. If PCR were routinely performed on all samples with DFA, it may improve sensitivity and increase the ability of laboratories to detect rabies cases with extremely sparse and non-uniform antigen distribution.

### Potential early rabies infection in two raccoon cases

Taken together, the low rabies virus level and observed distribution pattern (highest in the most caudal brain regions and lowest in rostral regions) are suggestive of early rabies virus infection. In laboratory animals, early infections are characterized by decreasing viral load from the brain stem to the forebrain, especially during peripheral, non-mucosal infections (27, 28), which is very similar to what was observed in these two PA raccoon cases. It remains unclear if animals are capable of transmitting virus during very early infections. For rabies virus to be transmitted, it must travel to the central nervous system and then back out to the periphery, specifically to the nerve endings in the salivary glands. Once in the salivary glands, rabies virus is secreted in saliva and can be transmitted by a bite. It would be interesting to see if virus was present in the salivary glands of these animals despite the low level of antigen and RNA in the brain; however, tissue was not available.

The presence of rabies virus neutralizing antibodies can interfere with infection and lead to low viral levels in the brain, which could explain the low antigen and RNA levels observed. Animals can develop virus neutralizing antibodies after vaccination, and vaccinated animals with sub-protective immunity can succumb to rabies virus infection (29–36). Oral rabies vaccination baits are distributed in western Pennsylvania as part of USDA’s raccoon rabies control program. The 2019 juvenile raccoon case was from Venango County in western PA, adjacent to the oral vaccination zone. It is possible that the 2019 juvenile raccoon was partially immunized but not fully protected from rabies infection, possibly through inherited maternal antibodies. The 2017 adult raccoon was collected in Carbon County in eastern Pennsylvania; it is unlikely this raccoon encountered oral vaccine. However, even in rabies enzootic and epizootic areas without wildlife vaccination, wild animals have been shown to have neutralizing antibodies attributed to acquired immunity from sublethal exposures (35, 37–42). Unfortunately, serum samples were not available from either animal for testing.

In this study, brain stem and rostral spinal cord were the most reliable tissue for rabies detection. Both DFA and PCR tests on cerebellum and hippocampus produced negative or indeterminate results for at least some replicates. The brain stem is one of the first brain structures where rabies virus is observed in natural infections or after experimental inoculation in peripheral muscle or foot (43–49). The increased reliability of brain stem and cerebellum for rabies diagnosis has been well documented in the literature, and insufficient sampling can lead to false negative results (43, 47, 50–54). For DFA, typical rabies antigen in the hippocampus and cerebellum can be more obvious due to large inclusions sometimes observed in pyramidal and Purkinje neuron somas (49). In early infections, antigen may present as dust-like particles in the axon bundles of the brain stem; although, more frequently inclusions of all sizes are also present. DFA testing personnel should be familiarized with both presentation types. A full cross section of brain stem and tissue from cerebellum or hippocampus is currently recommended for rabies diagnostic testing by WHO, OIE, and the US minimal national standard protocol (11–13, 50); spinal cord is not recommended for rabies diagnostic testing. It should be emphasized that neither DFA nor PCR can rule-out rabies if required brain areas are not available or recognizable.

### Investigation into potential cross-contamination

Cross-contamination can occur at several steps of tissue processing, sample preparation or during testing. An extensive search into potential contamination was performed for the 2019 juvenile raccoon case. Because the most likely source of contamination was positive samples processed around the same time, all such samples were sequenced. Sequencing was able to rule-out contamination because sequences from the juvenile raccoon displayed >1% differences to sequences from all positive samples processed at PABOL within two weeks of when the juvenile raccoon was processed.

## Conclusion

Accurate and timely primary diagnosis of rabies in animals is essential for subsequent post-exposure prophylaxis of exposed individuals. The 2017 and 2019 PA rabies cases demonstrate the sensitivity and objectivity of PCR in the identification of cases with low rabies virus as well as the need to test a cross section of brain stem for rabies diagnosis. These cases also highlight the importance of sampling, following standardized protocols, using multiple highly sensitive tests routinely, and submitting all samples with unexpected or atypical results to a reference laboratory for confirmatory testing especially when of public health importance.

## Acknowledgements

The authors would like to thank the additional members of the CDC Poxvirus and Rabies Branch and PABOL for help in planning, procuring funding, and discussion related to this project. This research received no specific grant from any funding agency in the public, commercial, or not-for-profit sectors. C.M.G. was supported in part by an appointment to the Research Participation Program at CDC, administered by the Oak Ridge Institute for Science and Education through an interagency agreement between the U.S. Department of Energy and CDC. The funders had no role in study design, data collection and interpretation, or the decision to submit the work for publication. Use of trade names and commercial sources is for identification only and does not imply endorsement by the Centers for Disease Control and Prevention, the Public Health Service, or the U.S. Department of Health and Human Services. The conclusions, findings, and opinions expressed by authors do not necessarily reflect the official position of the U.S. Department of Health and Human Services, the Public Health Service, the Centers for Disease Control and Prevention, or the authors' affiliated institutions.

